# A circuit model for transsaccadic space updating and mislocalization

**DOI:** 10.1101/2024.10.27.620527

**Authors:** Xiao Wang, Sophia Tsien, Michael E. Goldberg, Mingsha Zhang, Ning Qian

## Abstract

We perceive a stable, continuous world despite drastic changes of retinal images across saccades. However, while *persistent* objects in daily life appear stable across saccades, stimuli *flashed* around saccades can be grossly mislocalized. We address this puzzle with our recently proposed circuit model for perisaccadic receptive-field (RF) remapping in LIP and FEF. The model uses center/surround connections to store a relevant stimulus’ retinal location in memory as a population activity. This activity profile is updated across each saccade by directional connections gated by the corollary discharge (CD) of the saccade command. The updating is a continuous backward (against the saccade) shift of the population activity (equivalent to continuous forward remapping of the RFs), whose cumulative effect across the saccade is a subtraction of the saccade vector. The model explains forward and backward translational mislocalization for stimuli flashed around the saccade onset and offset, respectively, as insufficient and unnecessary cumulative updating after the saccade, caused by the sluggish CD time course and visual response latency. We confirm the model prediction that for perisaccadic RFs measured with flashes before the saccades, the final forward remapping magnitudes after the saccades are smaller for later flashes. We discuss the possibility that compressive mislocalization results from a brief reduction of attentional remapping and repulsion. Although many models of RF remapping, transsaccadic updating, and perisaccadic mislocalization have been proposed, our work unifies them into a single circuit mechanism and suggests that the brain uses “unaware” decoders which do not distinguish between different origins of neurons’ activities.

## Introduction

We make several saccades per second to foveate on different parts of a scene for high-resolution processing. Across a saccade the retinal image changes drastically, yet the world appears stable and continuous to us. Two main mechanisms have been proposed to explain this phenomenon of transsaccadic visual stability (TSVS): (1) the brain combines eye-position signals and retinotopic inputs to construct craniotopic (head-centered) representations (Andersen et al., 1985; Zipser and Andersen, 1988; Duhamel et al., 1997; Yang et al., 2024), and (2) the brain uses corollary discharges (CDs) of saccade commands to “compensate” for saccade-induced retinal changes (von Helmholtz, 1928; Duhamel et al., 1992; Wang et al., 2024). These mechanisms appear to contribute to transsaccadic space perception at long- and short-time scales, respectively (Poletti et al., 2013; Rutler et al., 2022). Here we focus on the CD mechanism because we would like to link its detailed, transsaccadic operations to the short-time-scale phenomenon of perisaccadic perceptual mislocalization. We consider saccades under the head-fixed condition so that the display screen for stimuli is craniotopic.

The original proposal of the CD mechanism is that the CD of a saccade cancels the retinal image motion produced by the saccade (von Helmholtz, 1928). A related observation is saccadic suppression: during saccades, visual perception (particularly of magnocellular stimuli such as motion) is impaired (Burr et al., 1994), and correspondingly, some visual neurons have reduced responses or reversed directional tuning (Richmond and Wurtz, 1980; Thiele et al., 2002). There is evidence that CDs are responsible for saccadic suppression (Richmond and Wurtz, 1980). However, although cancellation and/or suppression of saccade-induced retinal motion may contribute to TSVS, they are insufficient. Consider the double-step memory saccade task, a standard demonstration of TSVS, in which subjects sequentially saccade to two successively flashed and disappeared targets (Fig. 1). Since the first saccade (the rightward black arrow) changes the retinal position of the second target (from the magenta to green arrow), the brain must update the retinal position of the second target, by subtracting the saccade vector, before making the second saccade. Cancelling or suppressing the saccade-induced retinal motion would not provide the required updating. Moreover, the two saccades of this task can be made in total darkness; in this case there is no retinal motion to cancel or suppress but to make the second saccade, the brain still must update the retinal location of the second target.

**Fig. 1.**
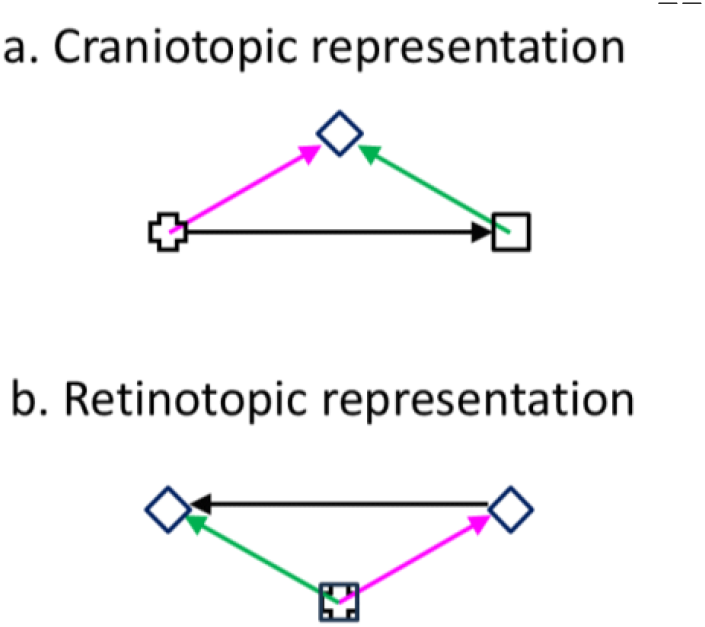
Double-step memory saccade task: the updating of the second target across the first saccade. (a) Craniotopic (screen) representation. After subjects fixate on the cross, the cross disappears, and the square and diamond are flashed successively. Subjects then sequentially saccade to the remembered square and diamond positions. (b) Retinotopic representation across the first saccade (back projected from the retina to the screen for comparison with a). The cross and square are superimposed as they correspond to the same retinal position, the fovea. The magenta and green arrows indicate the diamond’s retinotopic positions before and after the first saccade (the rightward black arrow in a), respectively. The leftward black arrow indicates that the diamond’s retinotopic position needs to be updated backward by subtracting the saccade vector.

The discovery of the CD-driven receptive-field (RF) remapping in LIP and FEF (Duhamel et al., 1992; Umeno and Goldberg, 1997; Sommer and Wurtz, 2006; Wang et al., 2016; So and Shadlen, 2022; Wang et al., 2024) has inspired new proposals on how the CD mechanism enables TSVS. The remapping refers to the observation that around the time of a saccade, cells’ RFs (perisaccadic RFs or pRFs) shift in the saccade (forward) direction. Early remapping studies focused on the fact that some cells show visual responses at their future (post-saccadic) RF (fRF) locations, accompanied by reduced responses at the current (pre-saccadic) RF (cRF) locations, even before the saccade onset. (A cell’s cRF and fRF are just its ordinary RF well before and well after the saccade, respectively; for a retinotopic cell, its cRF and fRF are offset by the saccade size on the display screen but superimpose on the retina. We use their screen (craniotopic) positions unless noted otherwise.) This leads to the Preview Theory of TSVS (Duhamel et al., 1992; Crapse and Sommer, 2012): On the screen, a cell’s fRF before a saccade will become its actual RF after the saccade. The activation of a cell by a stimulus in its fRF can thus be considered as giving the cell a preview of what will be in its RF after the saccade. Then, a comparison between the preview response and the postsaccadic (reafference) response can determine whether the world is stable or changed across the saccade.

Although intuitively appealing, the Preview Theory has a few difficulties. First, it requires cells whose pRFs remap completely to their fRFs without responses at their cRFs (or any other positions) before saccades. Otherwise, the preview responses would represent a mixture of stimuli in both the cRFs and fRFs, complicating the post-saccadic comparison. Second, the theory requires a downstream stage that stores the preview responses in memory and then compares them with the post-saccadic responses later. This memory and comparison stages have not been identified (Wurtz et al., 2011). Finally, for the double-step memory saccade task mentioned above, the flashed targets disappear before the first saccade, and they do not reappear to generate post-saccadic responses for comparison with the preview responses.

Later remapping studies revealed the details of the remapping time course in LIP and FEF (Wang et al., 2016; Wang et al., 2024). Although some cells respond to stimuli in their fRFs before the saccades, on average cells’ pRFs shift progressively from their cRF locations to near their fRF locations over time (from about100 ms before the saccade to about 100 ms after the saccade). The pRFs thus move through intermediate locations instead of jumping from the cRFs to the fRFs directly, posing further difficulties for the Preview Theory. There is, however, an alternative solution for TSVS (Wang et al., 2024). The progressive *forward* shift of pRFs from the cRF to fRF locations is equivalent to a progressive *backward* shift of the corresponding population response over the same time and distance (the saccade size) if the response is always considered a function of each cell’s cRF center position (i.e., the brain uses “unaware” positional decoders which always interpret a cell’s response as evidence for a stimulus in its cRF regardless of whether the response is indeed from the cRF stimulation or remapped from elsewhere (Qian et al., 2023); see Discussion). This backward shift of the population response effectively subtracts the saccade vector from a stimulus’ pre-saccadic retinal position to produce its correct post-saccadic retinal position (Fig. 1b).

The entire remapping time course must be driven by CDs because the stimuli for measuring the pRFs are flashed (and disappeared) before the saccade onset and there is no additional reafferent contributions to the pRFs during or after the saccade (Wang et al., 2024). This implies that the entire pRF remapping time course, including the portion after the saccade, can be viewed as predictive, and that what is remapped is the memory representations of the flashed stimuli. Then, to implement the above updating theory in a circuit model, we need a set of connections to maintain in memory the population response representing the retinotopic position of a flashed stimulus, and another set of connections, gated by the CD of a saccade, to shift the population response, across the saccade, to the updated position. We proposed the required connectivity patterns when modeling RF remapping in LIP and FEF (Wang et al., 2024). There are actually two types of RF remapping: the forward (or predictive) remapping discussed above and attentional (or convergent/compressive) remapping which is RF shifts toward attentional loci such as the saccade target (Connor et al., 1997; Zirnsak et al., 2014; Neupane et al., 2016; Wang et al., 2024). Inspired by related models for orientation-tuning dynamics (Teich and Qian, 2003; Teich and Qian, 2010), we explained attentional remapping with symmetric, center/surround connections among cells tuned to different retinotopic locations (red curve of Fig. 2a). This so-called Mexican-hat connectivity pattern is consistent with interactions among cells in LIP (Falkner et al., 2010) and FEF (Schall et al., 1995), and is also known to provide attractor dynamics for maintaining responses in memory (Cueva et al., 2021). We explained forward remapping with CD-gated directional connections (blue curve of Fig. 2a) that propagate responses backward from cells’ fRFs to their cRFs (Wang et al., 2016; Wang et al., 2024). These two sets of connections form a complete circuit for transsaccadic space updating to achieve TSVS (Zhang, 1996; Wang et al., 2024). Fig. 2b illustrates the updating of the second target across the first saccade of the double-step task (Fig. 1). The circles represent different cells’ cRF centers (in retinotopic coordinates). The second target (diamond) was flashed at the magenta cell’s cRF center, evoking a population response among the nearby cells (the red curve above the magenta cell) which is sustained by the center/surround connections (not shown in Fig. 2b) as a memory. Across the first saccade, this response profile is continuously shifted backward by the CD-gated connections (blue lines in Fig. 2b) to become a population response among the cells around the green cell (the red curve above the green cell), representing the updated retinotopic position of the second target. The total shift accumulated over time is equivalent to a subtraction of the saccade vector.

**Fig. 2.**
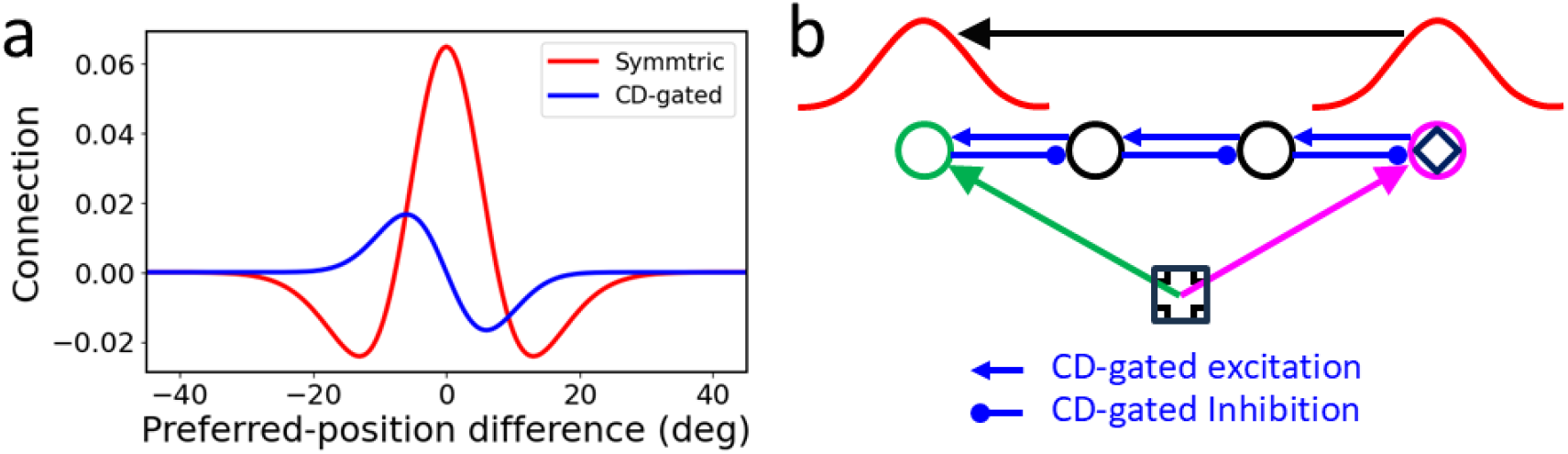
A circuit model for RF remapping and population-response updating across saccades. (a) Recurrent connection strengths among model LIP/FEF cells as a function of the difference between the cells’ preferred retinotopic positions (cRF centers). Symmetric, center/surround connections (red) can be modulated by attention to produce convergent remapping and directional connections (blue, for rightward saccades) are gated by CDs to produce forward remapping (Wang et al., 2024). (b) Schematic of the backward updating of the second target (diamond) across the first saccade of the double-step task of Fig. 1. Circles indicate a few cells’ cRF center locations in retinotopic coordinates. The diamond is flashed at the cRF center of the magenta cell, evoking a population response among nearby cells (red curve above the magenta cell) which is sustained by the symmetric connections (not shown) as a memory. This population memory response is shifted backward (black arrow) by the CD-gated connections (blue lines) across the saccade, updating the diamond’s retinotopic position from the magenta arrow to the green arrow. Note that on the screen, the green cell’s fRF is at the magenta cell’s cRF for the first saccade; the green cell will be activated by flashes at positions from its cRF to fRF with progressively longer delays, as observed in the forward remapping time course (Wang et al., 2016; Wang et al., 2024).

We also showed that the same circuit can update retinotpic positions of *persistent* stimuli across saccades (Wang et al., 2024). Although persistent objects in daily life appear stable across saccades, we mislocalize brief stimuli flashed around saccades, relative to those flashed well before or after the saccades, a phenomenon known as perisaccadic perceptual mislocalization (Matin and Pearce, 1965; Honda, 1991; Schlag and Schlag-Rey, 2002). The errors can be as large as many degrees of visual angle. If such errors occurred in daily life, our perception would be disturbingly unstable as objects would appear displaced after each saccade and then return to their correct positions when reafferent retinal inputs reach perception. Perisaccadic mislocalization has two components, a translational (or shift) component along the saccade axis and a convergent (or compressive) component toward the saccade target (Honda, 1991; Ross et al., 1997). The convergent component is smaller and larger, respectively, in the absence and presence of a postsaccadic visual reference, such as a ruler (Lappe et al., 2000). The translational mislocalization is in the saccade direction (forward) around the saccade onset, and disappears, or sometimes reverses the direction (backward), around the saccade offset (Honda, 1991; Lappe et al., 2000; Schlag and Schlag-Rey, 2002).

We argued previously that RF remapping alone cannot explain the observed mislocalization (Qian et al., 2023). We now demonstrate that under additional and reasonable assumptions, our circuit model of RF remapping that correctly updates persistent stimuli (and similarly, stimuli flashed well before or after saccades) for TSVS will produce the observed translational mislocalization for stimuli flashed around saccades. We focus on translational mislocalization because we interpret convergent mislocalization as reduced attentional repulsion relative to the baseline, a process distinct from transsaccadic updating and TSVS (see Discussion). The model makes testable predictions, and we confirmed one of them by reanalyzing our previous single-unit data from LIP and FEF (Wang et al., 2024). Our work clarifies, at the circuit level, the relationships between the physiological properties of RF remapping, the functional requirement of transsaccadic space updating, and the psychophysical observations of perisaccadic mislocalization, with implications on the nature of positional decoders used in the brain. The work suggests that translational mislocalization is really postsaccadic memory mislocalization of perisaccadically flashed stimuli.(Qian et al., 2023)

## Results

We consider a typical paradigm for perisaccadic perceptional mislocalization: A horizontal 12°saccade is made from an initial fixation point to a target (−6°and +6°relative to the screen center, respectively) while a probe stimulus is flashed at various times relative to the saccade onset; the location of the flashed stimulus is determined after the saccade. Fig. 1a can be reinterpreted as a configuration for measuring mislocalization, with the cross and square representing the initial fixation and target positions, respectively, and the diamond representing the flashed probe stimulus. For translational mislocalization the location of the flash does not matter; we assume the flash is at the screen center (0°) and its retinotopic position changes with the eye position at the time of the flash. For horizontal saccades, we need to consider only the horizontal spatial dimension in our simulations.

The circuit model consists of a one-dimensional array of LIP/FEF units representing the horizontal retinotopic space (Wang et al., 2024). The units receive feedforward inputs originated from the retina and are recurrently connected to receive lateral input from each other. A flashed spot on the retina can be viewed as a delta function in space and time. When this input reaches the recurrent, LIP/FEF units, we represented it as a Gaussian function in space and a gamma function in time to account for the intervening low-pass spatiotemporal filtering which produces spatial smear and temporal delay. The recurrent connections among the units are translationally invariant (Qian and Sejnowski, 1989) and can be divided into two sets. The first set follows a symmetric, center-excitation/surround-inhibition pattern among units tuned to different retinotopic positions (Fig. 2a, red curve). The second set is antisymmetric, directional connections gated by the CD of the saccade command with excitation and inhibition in the backward and forward directions, respectively (Fig. 2a, blue curve for rightward saccades). Since the physiological data show that forward RF remapping starts about 100 ms before the saccade onset and continues up to 100 ms after the saccade offset, we chose a similarly broad CD time course (Fig. 3,top row). The details of the model and its parameterization can be found in Methods; the model works with many different parameter combinations (Wang et al., 2024).

**Fig. 3.**
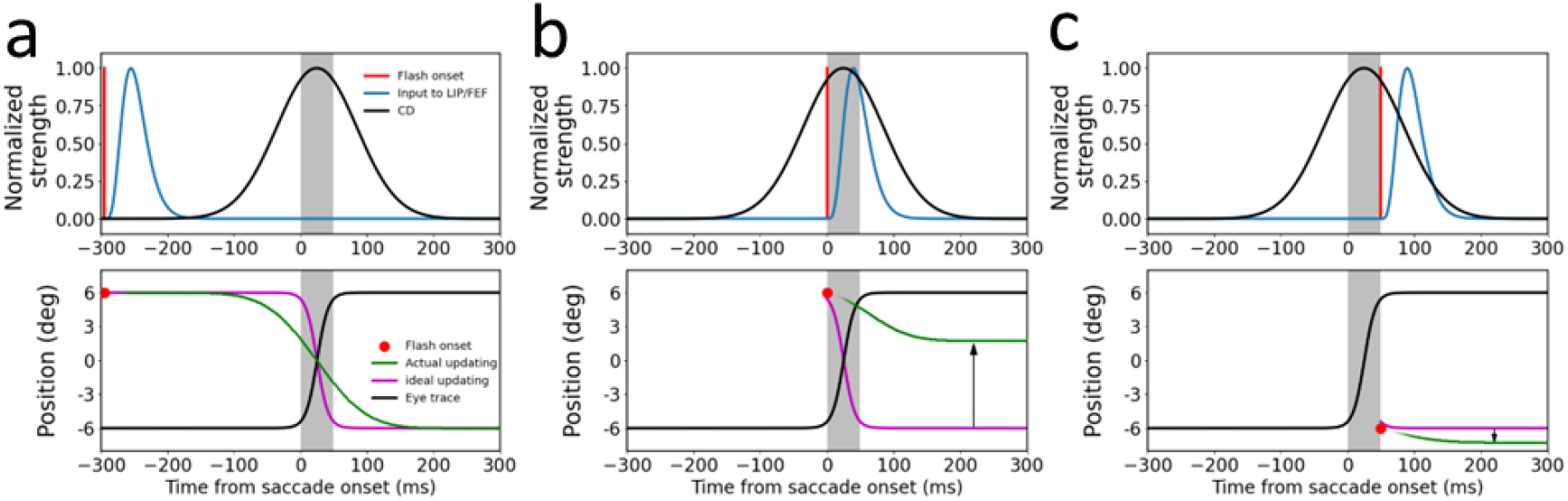
The circuit model explanation of translational mislocalization of stimuli flashed around saccades. The saccades are 12°rightward from -6°to +6°and the flashes are at 0°relative to the screen center. The gray shades indicate the 50-ms saccade duration. The three columns are the simulation results for the flash at (a) 295 ms before saccade onset, (b) saccade onset, and (c) saccade offset, respectively. The top row shows the temporal profiles of the flash on the retina (red), the input of the flash to the LIP/FEF units (blue), and the CD signal (black), with the peaks normalized to 1. The delay from the retinal flash to the peak of LIP/FEF input is 40 ms. The spatial profile of the input is a Gaussian (not shown). The bottom row shows the eye position (black) in the craniotopic coordinate (relative to the screen center), and the ideal (purple) and actual (green) updating of the flash’s position in the retinotopic coordinate (relative to the fovea/fixation). The ideal updating is simply the inversion of the eye position trace. The final differences (vertical arrows) between the cumulative actual and ideal updating after the saccade (any time after about 200 ms) is the mislocalization. The upward and downward arrows indicate forward and backward mislocalization for flashes at the saccade onset and offset, respectively.

We first considered the case when the stimulus is flashed 295 ms before the saccade onset (Fig. 3a). Despite the delay from the retina to LIP/FEF, the input (blue curve, top panel) reaches the LIP/FEF units before the start of the CD signal (black curve, top panel). This input is processed by the symmetric recurrent connections to produce a population response profile that stores the stimulus retinotopic position as a memory (Wang et al., 2024). We used the center-of-mass location of the response profile at a given time as the decoded retinotopic position of the stimulus at that time (green curve, top panel). When the saccade CD emerges, the memory response profile, and thus the decoded position, is updated backward by the CD-gated directional connections. We chose the CD strength such that the final, cumulative updating after the saccade is equal to the saccade size. Although the sluggish CD time course creates a mismatch between the ideal and the actual updating time courses, the finally updated retinotopic position, which stabilizes about 150 ms after the saccade offset (or 200 ms after the saccade onset), is accurate.

We next simulated how the same model responds to the stimulus flashed at the saccade onset (Fig. 3b). Because of the response delay from the retina to LIP/FEF and the CD signal starts before the saccade onset, by the time the input reaches the LIP/FEF units, it has missed much of the CD time course. Consequently, the cumulative backward updating of the memory response profile after the saccade is far short of the saccade size, resulting in a positional error in the forward direction (Fig. 3b).

We then considered the case when the stimulus is flashed at the saccade offset (Fig. 3c). Because the flash occurs when the eye has almost stopped moving, ideally there should be little updating of the retinotopic position of the stimulus. However, despite the response latency, the input to the LIP/FEF units still catches a tail part of the CD time course, and consequently the memory response profile is shifted backward slightly, producing a small positional error in the backward direction (Fig. 3c)

Fig. 4a summarizes the cumulative backward updating of the flash’s retinotopic position after the saccade as a function of the flash time relative to the saccade onset (green curve, top panel). Its difference from the ideal cumulative updating (black curve, top panel) is the mislocalization (black curve, bottom panel), which explains the translational component of the observed perisaccadic mislocalization. To explore the effect of the response latency from the retina to LIP/FEF, we shifted the gamma temporal response profile rightward by 20 ms so that the delay from retinal flash to the peak LIP/FEF input increases to 60 ms. As can be seen from the results in Fig. 4b,the forward and backward mislocalization of the flashes around saccade onset and offset becomes larger and smaller, respectively, with the longer input delay. This is expected because a longer input delay increases the missed portion of the CD time course which makes the backward updating of the flash around the saccade onset even more insufficient (i.e., larger forward mislocalization) and the unnecessary updating for the flash around the saccade offset smaller (i.e., smaller backward mislocalization). Conversely, if we reduce the response latency, or equivalently, if the CD profile is later than what we assumed in Fig. 3a,then the forward and backward mislocalization for the flashes around the saccade onset and offset will become smaller and larger, respectively (Fig. 3c). One way to manipulate the response latency is to change the stimulus contrast or size (see Discussion). Overall, the simulations are consistent with the observation that the forward mislocalization around the saccade onset is usually larger in magnitude, and more robust across studies, compared with the backward mislocalization around the saccade offset (Matin and Pearce, 1965; Lappe et al., 2000; Schlag and Schlag-Rey, 2002).

**Fig. 4.**
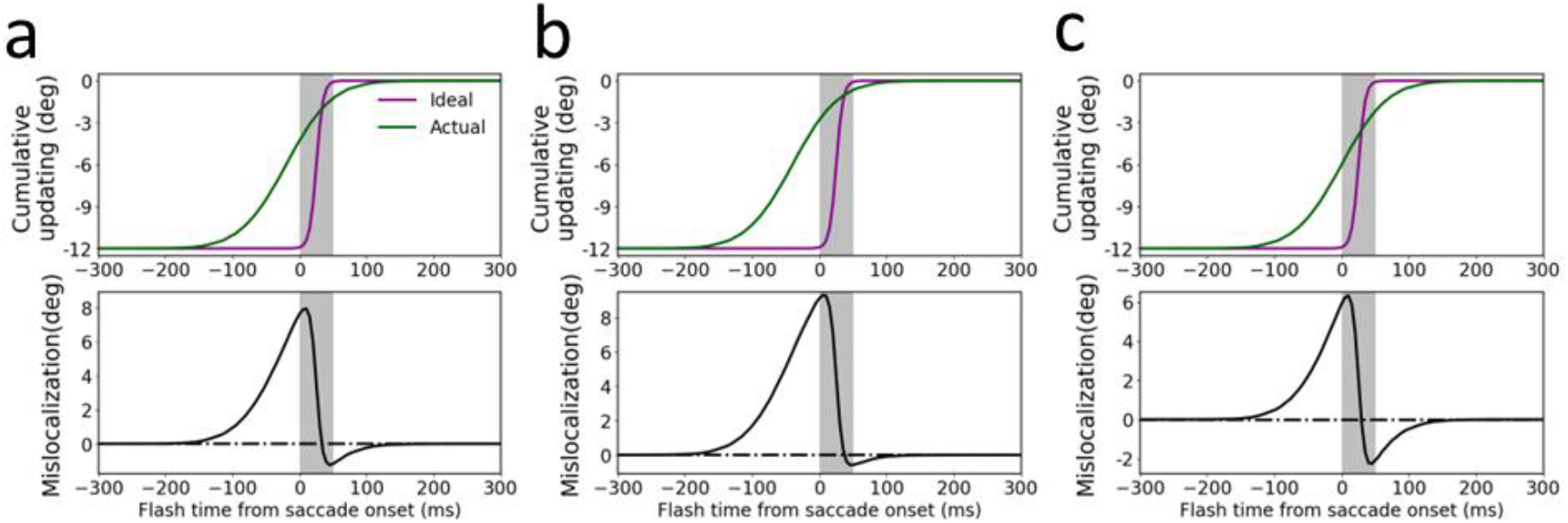
Post-saccadic cumulative updating and mislocalization for stimuli flashed at different times relative to the saccade onset. The three columns show results obtained with (a) the same parameters as in Fig. 3, (b) the delay from the retinal flash to the peak of LIP/FEF input increased by 20 ms (to 60 ms), and (c) the CD profile delayed by 20 ms. The top row shows the actual (green) and ideal (purple) cumulative updating of the flash’s retinotopic position after the saccade, and the bottom row shows their difference, the post-saccadic memory mislocalization. The ideal cumulative updating is the negative of the eye-position change from the time of the flash to the end of the saccade.

We finally apply the same model to a persistent stimulus and the result is shown in Fig. 5. The final, cumulative updating of its retinotopic position after the saccade is accurate, similar to the updating of the flash well before the saccade in Fig. 3a.

**Fig. 5.**
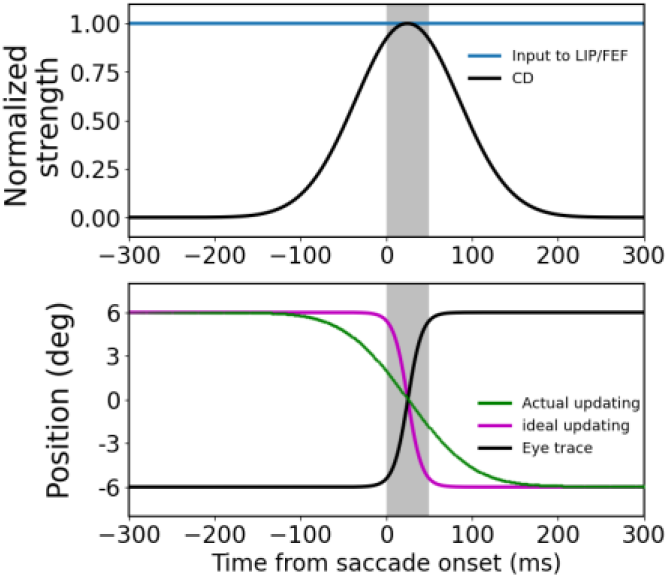
Transsaccadic updating of a persistent stimulus by the circuit model. The model parameters and the presentation format are the same as those for Fig. 3a except that the stimulus is always on at the screen center. The cumulative updating of the stimulus’ retinotopic position after the saccade (e.g., after 200 ms) is accurate.

Our model makes a few predictions (Discussion), one of which is the cumulative updating curves (green) in the top row of Fig. 4. They show that the later the flash, the smaller is the magnitude of the cumulative backward updating of the population response to the flash. It is particularly interesting to focus on flashes before the saccades because their retinotopic positions should all be updated backward by the saccade size (purple curves at -12°before 0 time) but the actual updating magnitude gets smaller as the flash gets closer to the saccade onset. This is equivalent to the prediction that for perisaccadic RFs measured with flashes before the saccades, the final remapping magnitudes after the saccades are smaller for later flashes. We reanalyzed our previously published single-unit data from LIP and FEF (Wang et al., 2024) to test this prediction. We first compiled the distributions of the perisaccadic flash time relative to the saccade onset for the LIP cells and FEF cells (Fig. 6d). For each brain area, we then divided a given cell’s trials into early and late groups according to the median time of the flash distributions (−100 ms and -113 ms relative to the saccade onset for LIP and FEF, respectively). Finally, we applied the same procedure as in (Wang et al., 2024) to determine the time course of the pRF remapping but for the early and late trials separately. Fig. 6a shows that the time courses of the forward remapping magnitudes in LIP and FEF; the mean remapping magnitudes are indeed greater for the early-flash trials (red) than for the late-flash trials (blue) in both LIP and FEF most of the time. Fig. 6b shows the mean remapping magnitudes from 160 to 260 ms after the saccade onset (or about 110 to 210 after the saccade offset). Although the differences between the early and late trials are small, they are significant (paired two-sided t-test, *t*_103_ = 2.39 and *t*_112_ = 2.30, and p = 0.019 and 0.023, for LIP and FEF, respectively). The small differences are expected because there were not many trials with the flashes close to the saccade onset (Fig. 6d). The physiological experiment was not designed for this test but we still find the predicted effect.

**Fig. 6.**
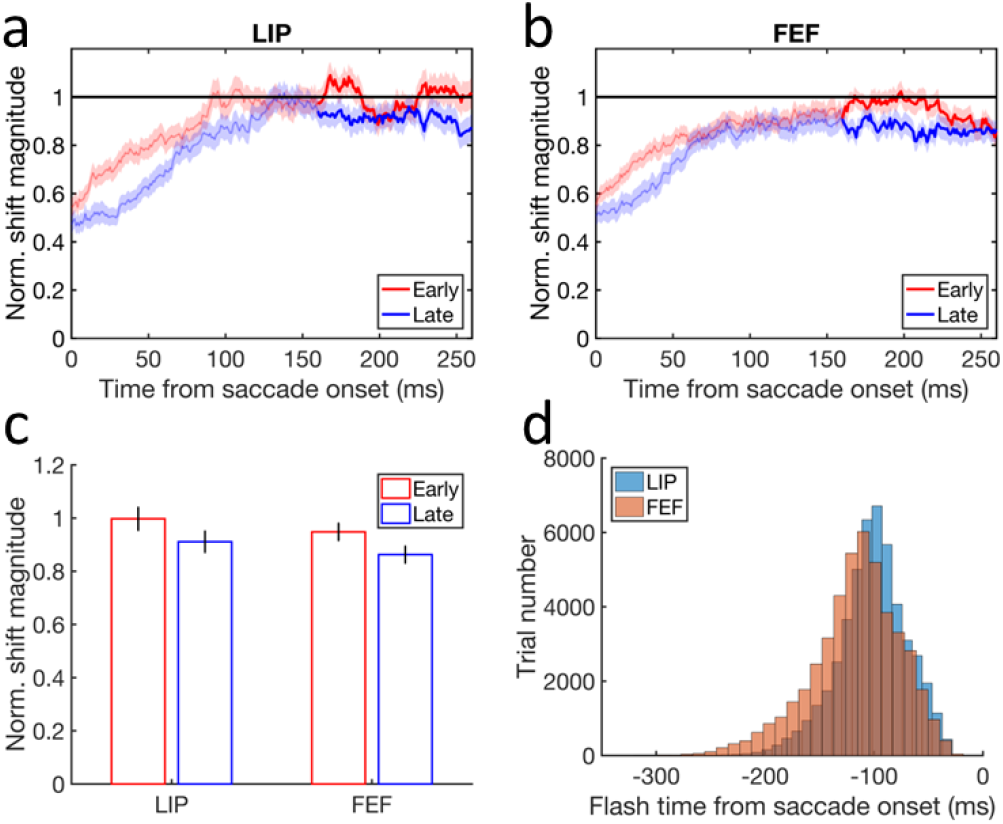
Testing the model prediction that for pRFs measured with flashes before the saccades, the final forward remapping magnitudes after the saccades are smaller for later flashes. (a-b) The time courses of the mean forward remapping magnitudes in LIP and FEF for the early (red) and late(blue) trials. The remapping magnitude is normalized by the corresponding saccade size before averaging over the cells. The horizontal line at 1 indicates a remapping magnitude equal to the saccade size. The shaded region around each mean curve indicates 1SEM. (c) The final mean forward remapping magnitudes from 160 to 260 ms after the saccade onset (highlighted portion in panels a and b) for the early (red) and late (blue) trials in LIP and FEF. The error bars indicate 1SEMs. (d) The distribution of the flash onset time relative to the saccade onset for the LIP and FEF cells. The numbers of the cells are n = 104 and 113 for LIP and FEF, respectively.

## Discussions

We argued previously that RF remapping *alone* cannot explain the observed perisaccadic perceptional mislocalization (Qian et al., 2023; Wang et al., 2024). First, when converting RF remapping into the corresponding population response for positional decoding, it is unclear whether the population response should be considered as a function of each cell’s remapped RF position (pRF center) or the original RF position before the remapping (cRF center). These choices imply that positional decoders are “aware” of and “unaware” of the remapping, respectively, and predict that the forward RF shift produces no shift and backward shift of the population response, respectively. Second, there is a mismatch between remapping studies and mislocalization studies: the former present perisaccadic stimuli before the saccade onset and measure RF shifts at different times across the saccade whereas the latter present a stimulus at different times across the saccade and measure its perceived position after the saccade. In this paper, we demonstrate that under some additional assumptions, which address the above two issues, the circuit model that uses CD-driven remapping/updating to achieve TSVS can explain the translational component of the observed mislocalization.

Our first assumption is that the forward remapping of retinotopic RFs is the sole mechanism for transsaccadic space representation over a few hundred ms around a saccade. This assumption is used in our simulations above as we decoded stimulus position solely from the updated retinotopic responses without considering craniotopic contributions. The assumption is consistent with the evidence that the CD-based remapping/updating mechanism and the eye-position-based craniotopic mechanism appear to operate at short (single saccades) and long (multiple saccades) time scales, respectively (Poletti et al., 2013; Rutler et al., 2022). The assumption also implies that the brain must use unaware positional decoders with which the forward RF remapping is equivalent to the backward shift (updating) of the corresponding population response to achieve TSVS (Qian et al., 2023). In contrast, with aware decoders, forward RF remapping is not equivalent to a backward shift of the population response and cannot be the mechanism for transsaccadic space updating.

To see the difference between aware and unaware decoders intuitively, consider the green cell in Fig. 2b which can be activated by either (1) the stimulation of its cRF or (2) the stimulation of the magenta cell’s cRF and then the lateral propagation of the activity from the magenta cell to the green cell via the CD-gated directional connections. For unaware decoders, the green cell’s activity is always evidence for a stimulus positioned at its cRF regardless of where that activity originates. For aware decoders, however, the green cell’s activity is evidence for a stimulus positioned at the green and magenta locations for the two cases, respectively. In other words, unaware decoders treat a cell as a fixed, labeled line whereas aware decoders “know” the origin of a cell’s activity and interpret it accordingly. Obviously, it would be much easier for the brain to implement unaware decoders than aware decoders but ultimately, this is an empirical issue to be settled by future experiments. Adaptation aftereffects provide indirect evidence for unaware decoders because aware decoders would “know” the adaptation-induced change of population responses and could null the aftereffects in principle (Xu et al., 2008; Seriès et al., 2009; Xu et al., 2012). Similarly, experiments showing perceptual effects of other types of RF dynamics support the use of unaware decoders in the brain (Gilbert, 1998; Fu et al., 2004).

Our second assumption is that in perceptual mislocalization experiments, the reported position of a flash is not decoded from the responses when they first reach LIP/FEF, but rather from the responses *after* the saccade, i.e., at the time of the report. For example, Fig. 3a shows that when a stimulus is flashed well before a saccade, the decoded position before the saccade onset would predict a backward mislocalization, but the decoded position *after* the saccade shows no mislocalization. Again, future studies are needed to evaluate this assumption. Our work suggests that perisaccadic perceptual mislocalization is actually postsaccadic memory mislocalization of perisaccadically flashed stimuli, lending further support to the notion that perceptual decoding often occurs in working memory (Ding et al., 2017; Luu et al., 2022).

Our final assumption is that persistent stimuli (and similarly, brief stimuli flashed well before saccades) are updated correctly across saccades without mislocalization (Teichert et al., 2010; Qian et al., 2023). This can be viewed as a definition of TSVS. Subjects may have individual biases in positional judgments unrelated to saccades, but those biases simply set the baseline against which perisaccadic mislocalization is determined. We thus did not consider such biases in our model.

In addition to the above assumptions, we also incorporated the following two facts into the model. First, the forward remapping in LIP and FEF has a sluggish time course that starts a little before the saccade onset and ends a little after the saccade offset (Wang et al., 2024). We assumed a correspondingly sluggish CD signal to drive the remapping/updating in the model. Second, there is a response latency from the retina to the remapping/updating stages such as LIP/FEF (Wang et al., 2016). We implemented the delay via low-pass temporal filtering and/or a hard time shift (Qian and Andersen, 1997). Because of the sluggish CD signal and the visual response latency, stimuli flashed at the saccade onset, whose retinotopic position should be updated backward by the saccade size after the saccade, would miss part of the CD time course, leading to insufficient backward updating and hence forward mislocalization. Stimuli flashed at the saccade offset, whose retinotopic position should not be updated, might still catch the tail of the CD time course, producing an unnecessary backward updating or backward mislocalization.

As we already noted, our model predicts that for pRFs measured with flashes before the saccade, the total forward remapping magnitudes after the saccade are larger for earlier flashes. We reanalyzed our previous single-unit data from LIP and FEF and confirmed this prediction (Fig. 6). This result also partially explains the observation that the final forward remapping magnitude after the saccade is a little smaller than the saccade size (Wang et al., 2024) as some of the flashes for measuring pRFs must have missed part of the CD time course. Another prediction of the model is that when the response delay from the retina to the stages of updating (LIP/FEF) is increased, the forward and backward translational mislocalization of flashes around the saccade onset and offset will become larger and smaller, respectively (Fig. 4). This could be tested by varying stimuli’s contrast and size: stimuli with greater contrast and size should have shorter response latency and therefore produce larger forward and smaller backward translational mislocalization of flashes around the saccade onset and offset, respectively. There is also a corresponding physiological prediction: for stimuli flashed at the same time right before the saccade, those with greater contrast and size should have larger final forward remapping magnitude after the saccade. Although our circuit model is one-dimensional and one directional, which is sufficient for simulating mislocalization during rightward saccades, it can easily be expanded to two spatial dimensions with different saccade directions (Wang et al., 2024).

Many ingredients of our model have been proposed previously but to our knowledge, they have never been integrated into a circuit model of RF remapping and TSVS to explain perisaccadic mislocalization. For example, early studies posit that during a saccade, the brain has a sluggish estimate of the eye position that first leads but then lags the actual eye position, producing forward and backward mislocalization around the saccade onset and offset, respectively (Matin and Pearce, 1965; Honda, 1991). Pola (2004) shows that when latency and persistence of visual responses to flashed stimuli are considered, a delayed but otherwise veridical eye-position estimate can account for the translational mislocalization. Teichert et al (2010) demonstrate that with physiological temporal filtering of visual inputs, the eye-position estimate that eliminates mislocalization for persistent stimuli produces translational mislocalization for flashed stimuli. Although we also include a sluggish signal (CD) and temporal filtering/delay, our model does not estimate the eye position but instead, updates stimuli’s retinotopic position, across saccades. More importantly, our model and the previous models assume that mislocalization arises from the stimulus memory *after* the saccade and the eye-position estimation *during* the saccade, respectively. Berreby and Krishna (2023) argue that anticipatory RF remapping can explain translational mislocalization. If their “Magnitude of forward remapping of the population response profile” (their Fig. 2AB) actually means our cumulative *backward* updating of the population response *after* the saccade, then their proposal and ours are conceptually similar. However, they directly drew the “remapping” curves in their Fig. 2AB whereas we mechanistically simulated the cumulative updating curves with our circuit model of TSVS. Our results cannot be derived from the forward RF remapping alone (Qian et al., 2023) but depend on the assumptions and facts discussed above.

While most studies found forward mislocalization for stimuli flashed around the saccade onset, two studies reported backward mislocalization instead (Jeffries et al., 2007; Weng et al., 2024). A key difference between the two studies and the rest is that the former provided veridical feedback of the flash position at the end of each trial whereas the latter did not. Why the feedback does not just eliminate or reduce the forward mislocalization but somehow overcompensates to produce the backward mislocalization is an open question. One possibility is that subjects might exaggerate the difference between the perceived stimulus position and the feedback position, as in many perceptual repulsion phenomena (Meng and Qian, 2005; Ding et al., 2017), which could lead to overcompensation through the feedback-driven learning.

We focused on the translational component of perisaccadic mislocalization. How, then, can the convergent or compressive component of the mislocalization be explained? We previously analyzed how various factors may affect convergent/divergent mislocalization (Qian et al., 2023), but if we assume that the brain uses unaware decoders, as we argued above, then we only need to consider attentional enhancement of responses around the saccade target, which produces attentional (convergent) RF remapping toward the target via the center/surround connections (Wang et al., 2024). [The notion that attentional remapping increases the cell density covering the attentional locus is only true under the aware-decoder assumption (Qian et al., 2023).] The attentional enhancement of responses alone “pulls” stimulus-evoked population responses toward the target whereas the attentional RF remapping alone “pushes” the population responses away from the target. Under physiologically reasonable parameters, the net prediction is a divergent mislocalization away from the target (Qian et al., 2023), consistent with the observed repulsion away from the attentional loci (Suzuki and Cavanagh, 1997; Pratt and Turk-Browne, 2003) and the enlargement of attended patterns (Anton-Erxleben et al., 2007). To explain convergent mislocalization of stimuli flashed around the saccade, which after a delay, activate LIP/FEF during and right after the saccade, we note that the attentional RF remapping toward the target in LIP and FEF starts to decrease about 50 ms before the saccade onset and is *diminished* during and right after the saccade (Wang et al., 2024), presumably because of reduced attention to the target over that period. Therefore, convergent mislocalization of stimuli flashed around a saccade might result from the diminished attentional remapping, and consequently diminished attentional repulsion, compared with the baseline before and after the saccade. Postsaccadic visual references increase convergent mislocalization (Lappe et al., 2000) perhaps by improving the perceived spatial relationship between the flashes and the target at the report time to reduce the smearing of the convergent pattern. Such smearing could affect convergent mislocalization more than translational mislocalization because the latter does not have the target as a convergent point.

Mislocalization of flashed stimuli similar to perisaccadic mislocalization has been produced by simulating saccade-like retinal motion but without the actual saccade (Ostendorf et al., 2006; Shim and Cavanagh, 2006). Such motion induced mislocalization of flashed stimuli in the absence of eye movements is known as the flash-lag effect (Brenner et al., 2006; Watanabe and Yokoi, 2006). This raises the possibility that perisaccadic mislocalization and the flash-lag effect might share similar underlying mechanisms (Teichert et al., 2010; Qian et al., 2023). Interestingly, motion can enhance lateral connections, in the motion direction, among cells tuned to different positions via spike timing dependent plasticity (Fu et al., 2004), similar to the CD gated lateral connections in our model. If the motion-enhanced connections are the mechanism for predictively updating the retinotopic positions of moving stimuli, then a circuit model similar to ours might explain the flash-lag effect. Future studies will hopefully clarify the relationships between different mislocalization phenomena and improve our understanding of neural mechanisms of space perception.

### Methods Circuit Model

We simulated a one-dimensional array of 360 LIP/FEF units covering 180° of horizontal retionotopic space, each unit governed by the equations:

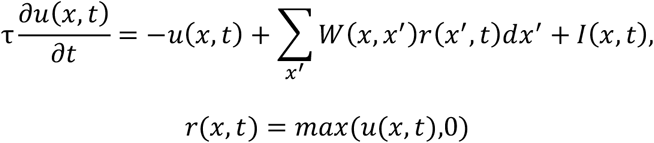

where *u*(*x, t*) and *r*(*x, t*) represent, respectively, the membrane potential and firing rate of the unit at location and time (*x, t*), *τ* is the membrane time constant, *W*(*x, x*′) is the recurrent connection strength from neuron at *x*′ to neuron at *x* and depends on (*x* − *x*′) only, and *I* is the feedforward inputs to LIP/FEF which originate from the retina. *W*(*x, x*′) is a sum of two parts: : (1) symmetric, center-surround connections modeled as a weighted difference between two Gaussians: *J*_*exc*_*G*(*x, x*′, σ_*exc*_) − *J*_*inh*_*G*(*x, x*′, σ_*ing*_) where 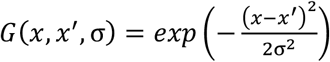, and (2) directional connections gated by the saccade CD, with the excitation and inhibition in the backward and forward directions of the saccade, respectively. For the simulations in this paper, we let *J*_*exc*_ = 0.165, σ_*exc*_ = 6°, *J*_*inh*_ = 0.1, σ_*inh*_ = 9.6°. For rightward saccades, we modeled the CD-gated connections as the antisymmetric, spatial derivative of the first Gaussian part of the center-surround connections: 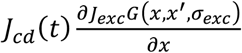 where the CD gating factor 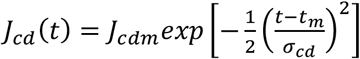 and *t*_*m*_ is the mid time of the saccade duration assumed to be 50 ms. For the simulations in Fig. 4c,we shifted *J*_*cd*_(*t*) t° the rightt by 20 ms. We let σ_*cd*_ = 60 ms, *J*_*cdm*_ = 0.97. The blue curve of Fig. 2a show the maximum directional connections when *t* = *t*_*m*_.

We considered both flashed and persistent visual inputs. A spot flashed on retina is filtered both spatially and temporally when it reaches LIP/FEF so we modeled its input to LIP/FEF units as a spatial Gaussian function and a temporal gamma function:

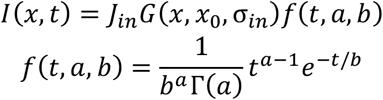

where *x*_0_ is the retinotopic position of the flash, and *a* and *b* are the shape and scale parameters, respectively. Translational mislocalization does not depend on the flash position. For the plots, we arbitrarily assumed a flash position of 0 in the screen coordinate; its retinotopic position varies with the eye position and is +6°and -6°before and after the 12 °saccade, respectively. We let σ_*in*_ = 4°, *J*_*in*_ = 2, *a* = 6, *b* = 8 ms so the delay from the retinal flash to the peak of the LIP/FEF input is (*a*-1)*b* = 40 ms. For the simulations in Fig. 4b,we added an additional delay of 20 ms by shifting the gamma function to the right by 20 ms so the total delay is 60 ms. We also considered a persistent stimulus turned on long before the the saccade onset and stayed on the screen center throughout. During the saccade, the Gaussian spatial profile of this input to the LIP/FEF units changes its retinotopic position according to the eye position and in the simulation (Fig. 5), this change is delayed by 40 ms to account for the visual response latency.

### Analysis of LIP and FEF single-unit data

We reanalyzed our LIP and FEF single-unit data in a published study (Wang et al., 2024) to test the prediction that for pRFs measured with flashes before the saccades, the final forward remapping magnitudes after the saccades are smaller for later flashes. The details of the experimental design and data collection and analysis can be found in that publication. Briefly, we recorded single units from monkeys’ LIP and FEF while they performed a delayed saccade task. For each unit, we measured its RFs from four different time periods (current, delay, perisaccadic, and future) by flashing a probe stimulus at one of the array locations in each period of each trial. For the current purpose, we focused on the cells’ RFs measured from the perisaccadic period (pRFs) and compared the remapping of the pRFs derived from the trials with early and late flashes. Specifically, we used the same 104 LIP cells and 113 FEF cells that passed our screening procedure under the saccade-onset alignment of repeated trials (Wang et al., 2024). We first compiled the distributions of the perisaccadic flash onset time relative to the saccade onset for all the LIP cells and all the FEF cells separately. Fig. 6d shows the results by dividing the time range of each brain area into 40 bins. For each area, we divided a given cell’s trials into early and late groups according to the median time of the flash distribution (−100 ms and - 113 ms relative to the saccade onset for LIP and FEF, respectively). Because of the relatively small number of trials at each flash location, the trials for some locations of some cells may all be placed in the early or late group. We used Matlab’s scatteredInterpolant function with the “natural” method to fill in the missing mean responses at those locations. We then applied the same procedure as in (Wang et al., 2024) to determine the time course of the pRF remapping but for the early and late halves of the trials separately (Fig. 6,a and b). We started the time-course plots at the saccade onset time (0) because that was when the pRF remapping directions in both LIP and FEF were mostly in the forward direction (Wang et al., 2024). Finally, we compared the final remapping magnitudes in the time window of 160 to 260 ms after the saccade onset (about 110 to 210 ms after the saccade offset) between the early-flash and late-flash pRFs (Fig. 6c).

## Acknowledgement

Supported by National Natural Science Foundation of China (32030045) and US National Eye Institute (R01 EY032938).

## Notes

https://data.mendeley.com/datasets/w6y53574zp/1

### Competing Interest Statement

The authors have declared no competing interest.

